# A cellular midbrain mechanism for executing fast and reliable escape

**DOI:** 10.64898/2026.07.01.734238

**Authors:** Yaara Lefler, Yu Lin Tan, Goncalo Ferreira, Alex Fudge, Marie Heffernan, Yeqing Wang, Tiago Branco

**Affiliations:** UCL Sainsbury Wellcome Centre for Neural Circuits and Behaviour, London, W1T 4JG, UK

## Abstract

Escape from threat is one of the most ancient and conserved sensorimotor transformations and must satisfy two competing demands. It must be fast and reliable, because failing to escape genuine threats risks death, yet also selective, because escaping indiscriminately costs energy and missed resources ^1,2^. Speed and reliability favour a system with a low threshold that responds to the slightest indication of danger, whereas selectivity demands a high threshold that filters out innocuous stimuli — yet both must be achieved simultaneously. While previous work has identified mechanisms for implementing selectivity ^3–6^, it is not known how the mammalian brain achieves speed and reliability once genuine threats have been identified. Here we use whole-cell recordings from dorsal periaqueductal gray (dPAG) neurons in behaving mice to show that the escape circuit resolves this tension via high intrinsic excitability at the single cell level. We find that while the membrane potential of dPAG neurons does not reach action potential threshold during exploratory behaviour, they require only a small amount of current to fire. Threat stimuli cause a sparse increase in synaptic input rate that, because of the high excitability, produces a sustained depolarizing voltage step that drives spiking and escape behaviour. Individual dPAG neurons respond similarly on escape and non-escape trials —what determines escape is the fraction of the dPAG population that is recruited, a finding we confirm with single-unit recordings in freely moving mice. The membrane voltage step response is also invariant to threat type, in contrast to superior colliculus neurons where membrane potential dynamics reflect stimulus identity. These findings reveal a single neuron mechanism for fast and reliable escape, in which the high intrinsic excitability of dPAG neurons transforms sparse, threat-evoked synaptic input into a rapid and stereotyped voltage signal, converting diverse threat representations into a uniform escape command.

## Main

Escape is a defensive locomotor action that moves an animal away from threat and towards safety, and which should be initiated as quickly and as reliably as possible once a genuine threat has been detected ^7,8^. In vertebrates, the periaqueductal gray (PAG) is the principal circuit for commanding escape initiation, receiving convergent threat information from multiple brain areas ^9–12^. A major source of this input is the superior colliculus (SC), whose deep layers contain neurons that are strongly activated by threatening stimuli and convey threat signals to the PAG through a direct excitatory projection ^3,13^. Previous work has shown that the SC–PAG connection is weak and unreliable and imposes a synaptic threshold on escape initiation, effectively filtering out inputs that do not generate sufficiently strong SC activity to drive PAG neurons ^3^. While this mechanism achieves selectivity, it is not clear how it is compatible with the fast and reliable PAG activation that escape demands, because the same synaptic bottleneck that filters weak inputs also constrains the excitatory drive available to PAG neurons during genuine threats. Understanding how PAG neurons resolve the tension between selectivity and reliability requires measuring synaptic integration during escape behaviour, which has not been technically possible.

### dPAG neurons are quiescent yet highly excitable

To access the subthreshold dynamics of PAG neurons during escape, we established a head fixed paradigm for whole-cell patch-clamp recordings while mice escape to a shelter. The setup consists of an air table and a translatable floating arena ^14–17^ with a shelter zone, allowing the mouse to navigate and explore the physical environment despite being head fixed (Fig. 1A; Extended Data Video 1). Exploratory patterns were similar to those of freely behaving mice in an open arena, with a strong preference for the shelter area (fraction of time moving: 16.4±7.9%; fraction of time in shelter: 34.3± 23.3%; N = 61 mice; Fig. 1B). Presentation of threatening sound stimuli elicited reliable escapes to shelter with high probability (65.5 ± 27.1%, N = 21 mice) and short reaction times (1.68 ± 1.15 s, n = 79 trials, N = 21 mice; Fig. 1C; Extended Data Video 2), with trajectories similar to those of freely behaving mice. We then adapted this setup to obtain stable whole-cell recordings from dorsal PAG neurons during escape (N=176 cells, N=98 mice; Extended Data Fig. 1; see Methods).

**Figure 1:**
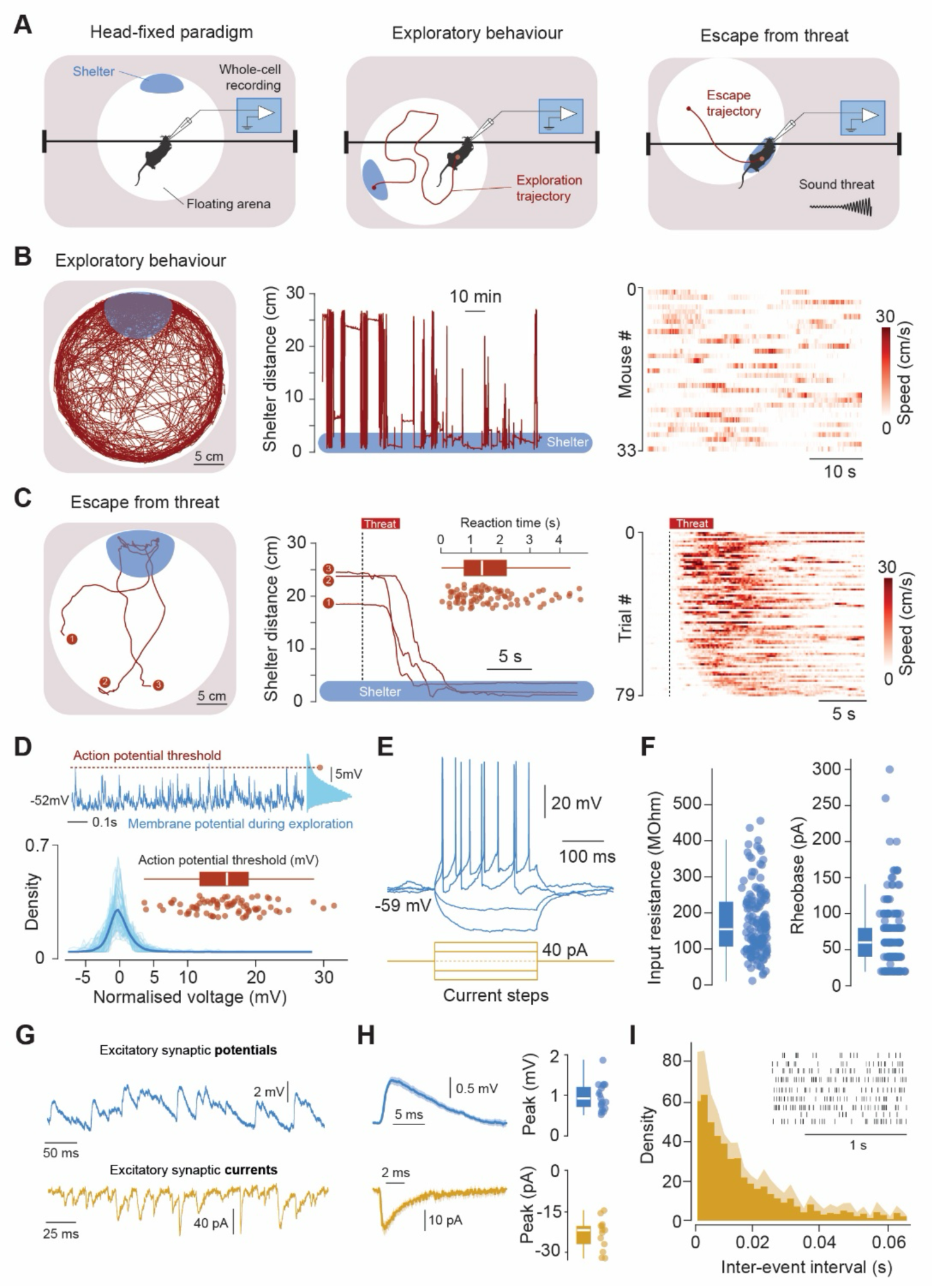
dPAG neurons are silent but highly electrically excitable. **A,** Left, illustration of the head-fixation setup, consisting of a floating and XV-translatable light-weight circular arena with a shelter zone. Illustration of exploration (middle) and threat-evoked escape trajectories (right). **B,** Left, exploration trajectory for a full experimental session (∼2.5h), and distance to shelter for a subset of this period (middle). Right, speed heatmaps of spontaneous locomotion at random times in the session (n=33 epochs, n=33 mice). **C,** Example of three escape trajectories following a sound threat, shown from running onset to escape termination (left), and corresponding distance to the shelter aligned to threat onset (middle). Inset, summary of escape reaction times (n=79 trials, n=21 mice; median 1.38s). Right, heatmaps of locomotion speed in escape trials (n=79, n=21 mice), sorted by reaction time. **D,** Top, example of resting membrane voltage (Vm) of a PAG cell and corresponding Vm histogram on the right. Dashed line shows the action potential (AP) threshold of that cell. Bottom, Vm distributions of each cell (n=77 cells; Vm has been normalised to the mean Vm). Inset, summary of AP thresholds (n=77 cells). **E,** Example response of a PAG cell to 250 ms step current injections. **F,** Summary of input resistance (left) and rheobase (right) for 154 cells. **G,** Example voltage (top) and current (bottom) recordings of synaptic input during exploratory behaviour. **H,** Average EPSP waveform and summary of peak amplitudes (top) and same for EPSCs (bottom). I, Histogram of inter-event intervals for spontaneous EPSCs, and example event rasters (inset, n=10 cells). Box-and-whisker plots show median, interquartile range and spread, with individual datapoints overlaid. Shaded area in histograms represents 95% confidence intervals. Shaded area around lines shows s.e.m.

We first measured the membrane potential during exploratory behaviour. We found that the membrane potential distribution was remarkably narrow (mean Vm: −50.01±9.15mV; mean s.d.: 1.5±0.7mV; N=84 cells) and very rarely crossed the action potential (AP) threshold (Fig. 1D). This is consistent with previous observations that dPAG neurons fire very sparsely outside escape ^3,18^, and with the requirement for selectivity: because dPAG spiking drives escape, spontaneous firing during exploration would trigger inappropriate flight responses. To understand the biophysical basis of this quiescent state, we measured intrinsic properties using somatic current injections and found that dPAG neurons have high input resistance (185.1 ± 96.1 MΩ, N = 156 cells), a membrane time constant of 17 ± 10.6 ms, and an action potential threshold 16.5 ± 6.4 mV above resting membrane potential (N = 166 cells; Fig. 1E-F). As a result, dorsal PAG cells have a very small rheobase (67.3 ± 46.6 pA), indicating high intrinsic excitability *in vivo*.

This finding is surprising because highly excitable neurons should be susceptible to large voltage fluctuations and spontaneous spiking, which would compromise the selectivity that their silence during exploration requires. We therefore took advantage of the high resolution of whole-cell recordings to characterise synaptic input directly. Excitatory postsynaptic potentials (EPSPs) had a unitary amplitude of 0.97±0.3 mV (N = 19 cells), and voltage-clamp recordings revealed a unitary current amplitude of −23±5.5 pA (N = 13 cells; Fig. 1G-H). While individual synaptic events are thus large relative to the voltage threshold and rheobase, the input rate during exploration was insufficient to drive spiking (mean inter-event interval = 0.21+0.052 s, mean frequency = 48.3 Hz; Fig. 1I), leaving dPAG simultaneously silent yet exquisitely poised to fire upon even a small increase in input.

### Threat input triggers sustained depolarizing voltage steps

We next presented auditory threat stimuli and measured membrane potential responses during escape. Threats caused a sharp and sustained step depolarization (Fig. 2A-B), with an average peak amplitude of 9.98±4.05 mV (N = 71 cells; depolarizing responses observed in 66.2% of recorded neurons). To determine the molecular identity of responsive neurons, in a subset of experiments we expressed ChR2 in VGluT2+ neurons and identified cell type using optotagging ^20^. While threat-evoked depolarizing responses were observed in seven of eight VGluT2+ neurons, none of the non-VGlutT2+ neurons depolarized (N = 3; Extended Data Fig 2), indicating that the threat-evoked voltage step occurs selectively in the excitatory dPAG population that commands escape actions ^3,18,20,21^.

**Figure 2:**
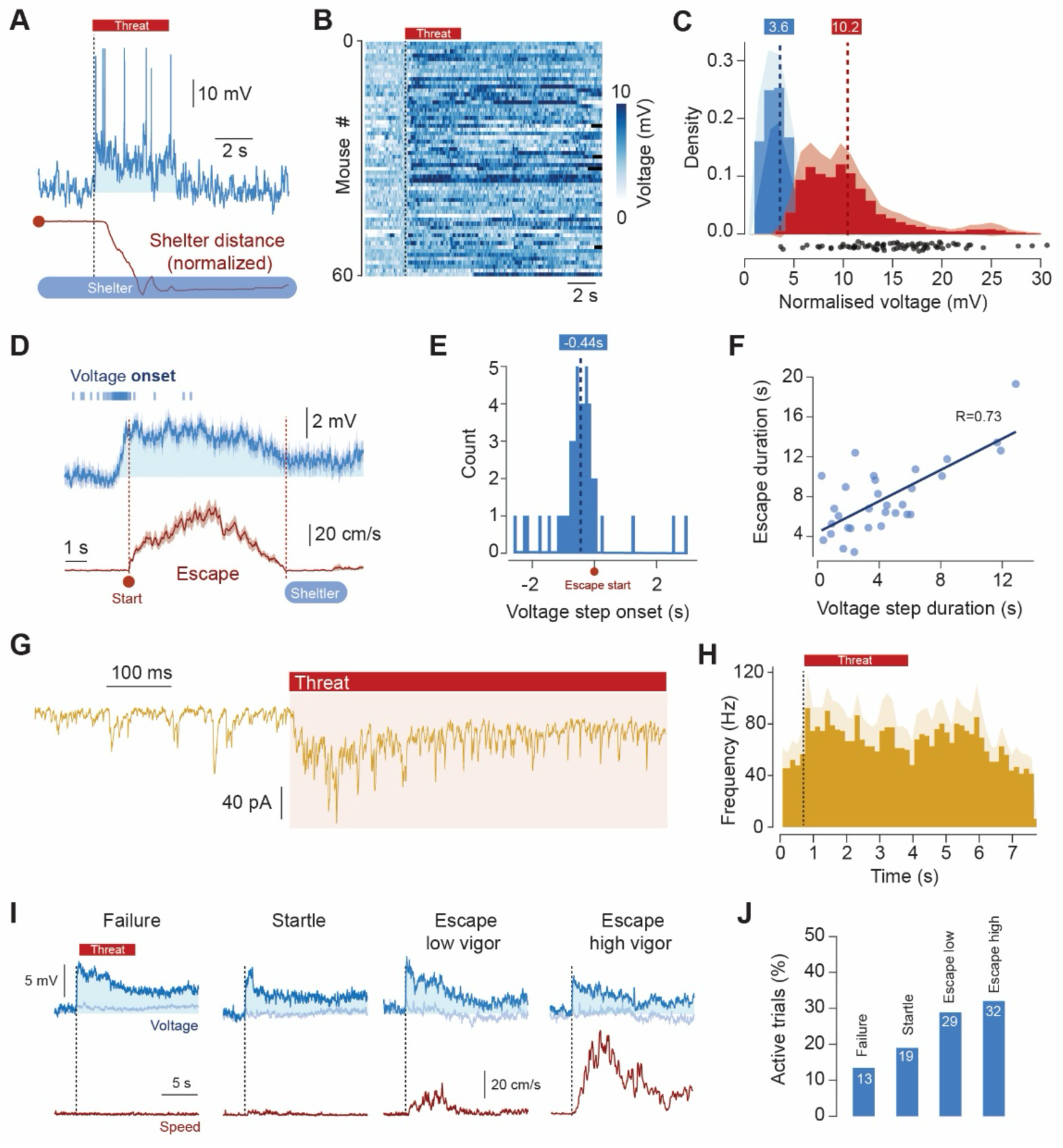
Threat evokes sustained voltage steps that drive escape. **A,** Example of escape trial showing the membrane voltage (Vm) of a PAG cell and corresponding mouse distance to shelter following a sound threat (dashed line marks threat onset). **B,** Vm heatmaps for high vigour escape trials (n=60 trials, n=43 cells; Vm is normalised to the mean Vm of each trial). **C,** Average distributions for the top 5% voltage values of mean normalised Vm during exploration (blue; mean=3.6 mV) and threat (red; median=10.2 mV). **D,** Time-warped average Vm (blue) and speed (red) during escape (n=50 trials, n=37 cells). Lines above the trace show onset of the voltage step for each trial. **E,** Distribution of voltage step onsets aligned to escape onset (median=0.44s before escape; n=50 trials). **F,** Voltage step duration versus escape duration (Pearson r = 0.73, P=3.85×10-□,n=31 trials, n=23 cells). **G,** Example trace of synaptic currents during threat. **H,** EPSC frequency during escape from threat (n=6 trials, n=6 cells). I, Average Vm (blue) and speed (red) aligned to threat onset (dashed line) for four behavioural outcomes (failure, n=164 trials; startle, n=79 trials; low-vigour escape, n=45 trials; high-vigour escape, n=50 trials). Voltage averages are separated into depolarizing (dark blue) and non-depolarizing trials (light blue). **J,** Fraction of active trials for each behavioural outcome, for the same trials shown in I. In all plots the shaded areas around lines show s.e.m; filled area under voltage traces shows periods above baseline. Shaded areas in histograms represent 95% confidence intervals.

A consequence of this step response was to bring the membrane potential closer to the action potential threshold (mean distance to AP threshold: 7.4±1.0 mV, significantly reduced from baseline, P = 1.1×10^−11^ Wilcoxon signed-rank test; Fig. 2C), increasing spiking probability 2.7-fold from the onset of the response and throughout its duration (P = 0, Wilcoxon signed-rank test, N = 179 trials from 97 cells). Voltage step onset preceded escape initiation by 0.44 ms, which coincided with the peak of the depolarization (Fig. 2D-E), and escape was sustained for the duration of the evoked voltage step (correlation between step duration and escape duration: r = 0.73, P = 3.85×10⁻⁶; Fig. 2F). These data show that threat input is converted into a voltage step that sustains dPAG escape neurons at spiking threshold.

To understand the synaptic basis of this sustained depolarization we recorded synaptic currents during threat presentation and escape. Input rate increased at threat onset to 74.6±16.9 Hz and remained elevated throughout (P=0.0047, paired t-test relative to baseline mean of 48.3±13.4; Fig. 2G-H). The increase from baseline was only ∼25 Hz, but because of the high intrinsic excitability of dPAG neurons, this small change in synaptic drive was sufficient to produce a large, sustained depolarization.

Since escape is probabilistic, we next took advantage of this variability to ask what distinguishes escape from non-escape trials at the level of single-neuron membrane potential dynamics. Surprisingly, we found that in neurons that responded, the depolarizing steps were indistinguishable between escape and non-escape trials (P = 0.78, t-test; Extended Data Fig. 3B). However, the fraction of neurons activated on non-escape trials was 56% lower than on escape trials, suggesting that escape initiation is determined not by the magnitude of depolarization in individual neurons but by the fraction of the population that is recruited (13.4% vs 30.5%; Extended Data Fig. 3B). We tested this further by comparing escapes of different vigour: the step amplitude was again identical across conditions (all comparisons t-tests p>0.1; N = fail 22/164; startle 15/79; low escape 13/45; high escape 16/50), and the difference was carried by the fraction of neurons that underwent the step (Figure 2I-J). In experiments where we recorded multiple trials from the same neuron, we confirmed that activation was probabilistic at the single-cell level — each neuron was more likely to undergo the step on escape trials than on non-escape trials (Extended Data Fig 3A).

### Escape depends on how many PAG neurons respond rather than how strongly

The intracellular data suggest a model in which threat input switches individual dPAG neurons from a quiescent resting state to a depolarized, spiking state, with little intermediate response, and that escape is initiated when a sufficient fraction of the population is switched on. To test this directly at the population level, we recorded single-unit activity from dPAG using chronically implanted Neuropixels probes in freely moving mice (Fig. 3A), allowing us to simultaneously sample many neurons and compute the fraction active on each trial.

**Figure 3:**
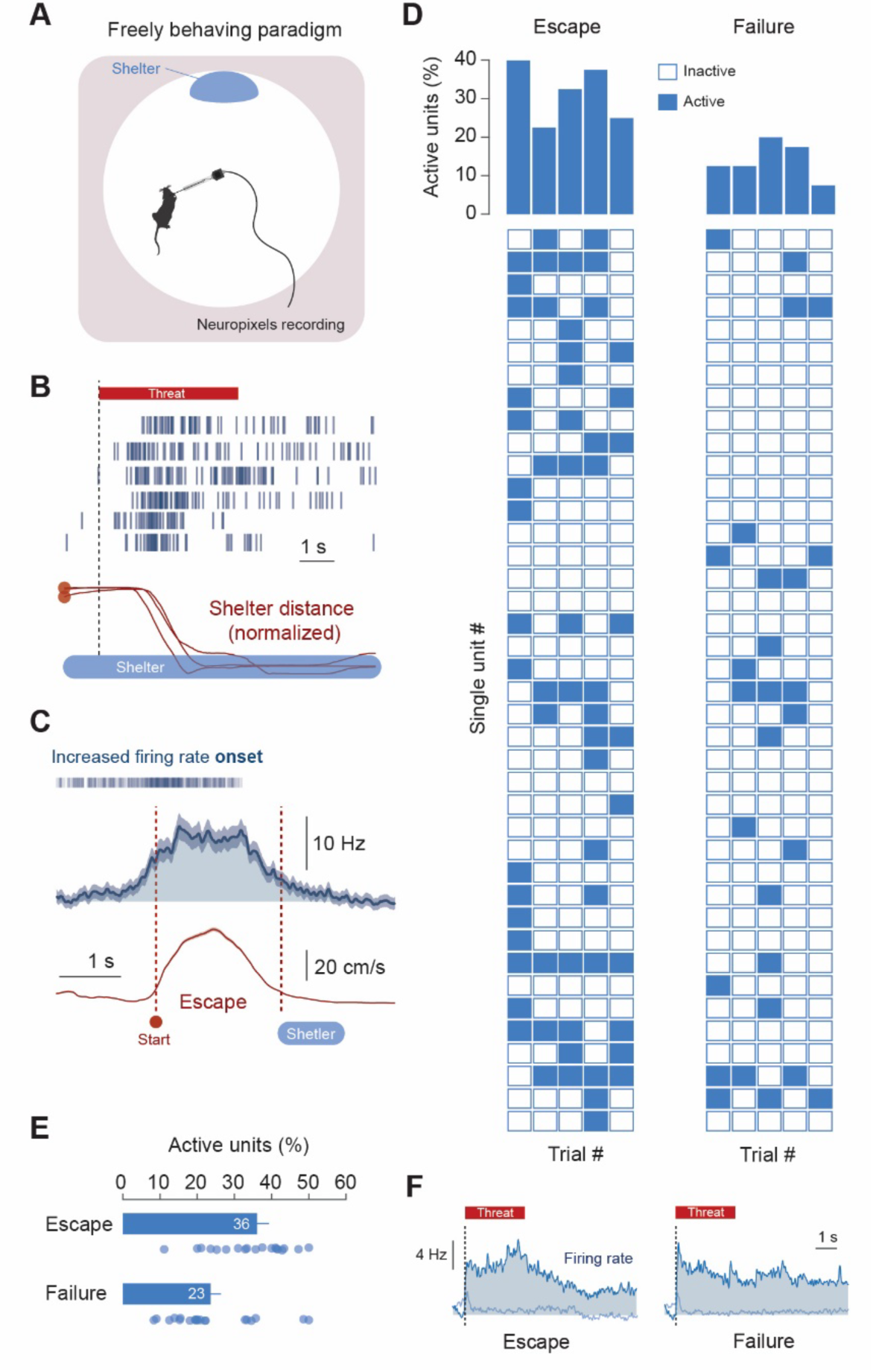
Escape depends on the fraction of active dPAG neurons. **A,** Illustration of the setup for freely moving mice, with an elevated circular arena (92 cm diameter) and a dark shelter. **B,** Top, example raster plots of spike times for escape trials following threat (n=3 trials and 2 units). Bottom, corresponding distance to shelter profiles for the three trials, aligned to threat onset. **C,** Time-warped average PSTH (blue) and speed (red) from escape onset to shelter entry (n=507 trials, n=331 units, n=5 mice). Lines above the PSTH trace shown the onset of firing rate increase for each trial. **D,** Bottom, grid plot of trials per unit, divided into escape (left) and failure trials (right), for an example session. Filled boxes show trials with a significant increase in firing rate. Top, average percentage of units that increase their firing rate for escape (left) and failure (right) trials, for this session. E, The average percentage of units per trial with increased firing rate in escape (blue) or failure (red) trials, for all sessions (n=22 sessions, 1615 escape trials, 2330 failure trials, 331 units, n=5 mice; paired t-test p=0.0006). **F,** Average threat-aligned PSTH of all units with increased firing rate in escape (dark blue left, n=507 trials) and failure trials (dark blue right, n=469 trials), and corresponding PSTHs for non-active units in light blue. In all plots the shaded area around lines are s.e.m; filled area under PSTH traces shows periods above baseline; bar plots show mean and s.e.m, with individual datapoints for each session overlaid.

In the experimental paradigm, mice freely explored an open arena with a shelter, and threat stimuli elicited rapid escapes with probability of 34.9±22%, reaction time of 2.1±2.1s and maximum speed of 63.4±26.4 cm/s, as previously reported ^22–24^. Consistent with the intracellular data, escapes were accompanied by a sharp increase in firing rate at escape onset in 76.0% of units (mean firing rate increase: 16.83±0.69 Hz (s.e.m); 507 trials from 259 units), and elevated firing was sustained throughout escape (Fig. 3B-C). We then identified the population of units that responded with increased firing rate on at least one trial, and for each trial quantified the fraction of this population that was significantly activated (Fig. 3F). The fraction of responsive neurons was ∼55% higher on escape than non-escape trials (22 sessions, 1,615 escape trials, 2,330 failure trials from 341 active units; P = 0.0006, paired t-test). Critically, the firing rate profiles of individual units that did respond were similar between escape and non-escape trials (P = 0.14; Mann-Whitney test, Fig. 3F), mirroring the invariance of the subthreshold voltage step observed in whole-cell recordings. These population data confirm that escape is determined not by how strongly individual dPAG neurons are activated, but by how many are recruited: threat input stochastically switches neurons into a high-firing state whose profile is relatively invariant, and escape occurs when the fraction of switched neurons is sufficiently large.

### PAG neurons transform diverse threat signals into a uniform escape command

Our experiments so far used a single type of threat stimulus. However, animals must escape from threats of different sensory modalities, and behavioural studies have shown that escape behaviour in mice is remarkably similar regardless of the stimulus that elicits them ^25–27^. The model emerging from our data offers a potential explanation for this invariance: if the high intrinsic excitability of dPAG neurons causes them to switch into a stereotyped active state in response to sufficient synaptic input, then the nature of the input should not matter — any threat signal that recruits enough synaptic drive should produce the same voltage step and the same escape output. To test this, we presented head-fixed mice with visual looming stimuli ^24,25^, which reliably elicited escapes to shelter with trajectories and reaction times comparable to those evoked by auditory threats (Fig. 4A-C). We then performed whole-cell recordings from both dPAG neurons and neurons in the deep layers of the superior colliculus (SC), the principal source of sensory threat information to the dPAG, and compared responses to visual and auditory threat stimuli by computing the mean squared difference between all pairs of responses within and across threat types. In SC neurons, membrane potential dynamics reflected the identity of the stimulus, with average pairwise differences significantly greater across threat types than within (within label: 24.4±1.2, across label: 37.3±2.3, P = 0.000002, Mann-Whitney test; Fig. 4C-D). In dPAG neurons, by contrast, both visual and auditory threats generated a sustained step depolarization, and the average pairwise difference was the same within and across threat types (within label: 31.5±0.8, across label: 31.8±0.8, P = 0.679, Mann-Whitney test; Fig. 4C-D). dPAG neurons therefore discard information about threat identity and respond with a stimulus-invariant voltage step. This demonstrates that the conversion from graded, stimulus-specific threat representations in the SC, to a uniform escape command occurs at the level of individual dPAG neurons: the biophysical properties that produce the all-or-none voltage step effectively standardise diverse inputs into a common escape currency, ensuring that the speed and reliability of escape are preserved regardless of how the threat was detected.

**Figure 4:**
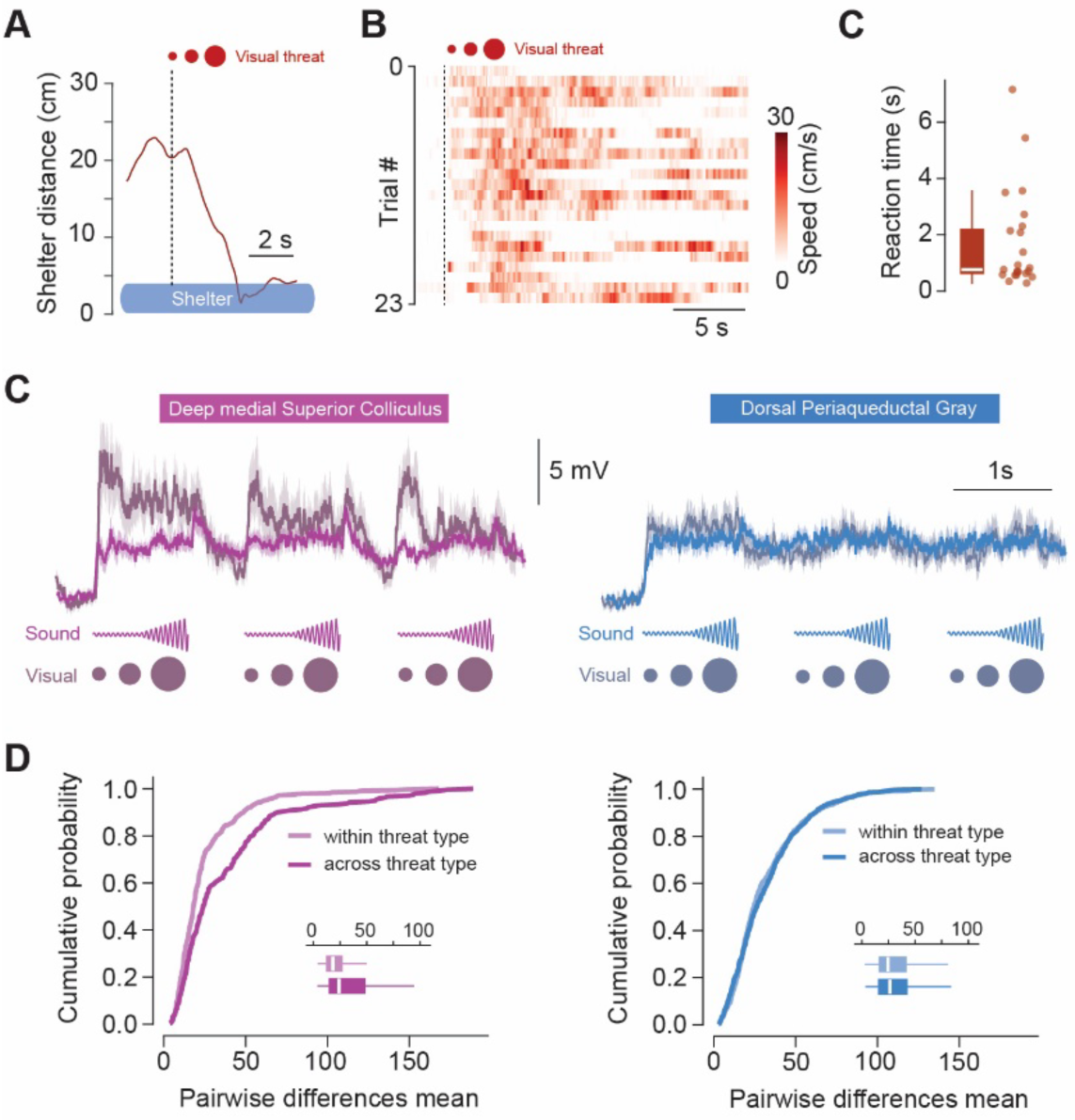
dPAG responses to threat are independent of stimulus identity. **A,** Left, distance to the shelter during an example escape response to visual threat, aligned to threat onset (dashed line). B, Speed heatmaps of escape trials (n=23 trials, n=14 mice), sorted by reaction times. C, Summary of escape reaction times (n=23 trials, n=14 mice; median 0.76 sec). C, Average membrane voltage (Vm) responses in superior colliculus (SC(cells (left) and PAG cells (right) for sound (n=37 PAG trials and n=24 SC trials) and visual (n=18 PAG trials and n=10 SC trials) threat stimuli (n=46 PAG cells and n=19 SC cells). D, Cumulative probability of the mean squared difference of Vm between trials, for within threat type and across sound and visual threats, for voltage responses in SC (left) and PAG (right). Insets box-and-whisker plots show median, interquartile range and spread, with individual trials’ data overlaid; shaded lines are s.e.m.

## Discussion

Escape must be simultaneously selective, fast, and reliable — demands that appear to be in tension because selectivity favours a high activation threshold while speed and reliability favour a low one. Our data suggest that the dPAG escape circuit resolves this tension by separating these demands across levels of organisation. While selectivity is achieved at the synaptic level through the unreliable SC–PAG connection that filters out inputs that do not generate strong presynaptic activity ^3^, speed and reliability are achieved at the single-neuron level: dPAG neurons are highly excitable, sit close to spike threshold, and require very little synaptic input to fire, such that input patterns that get through the SC-PAG synaptic bottleneck generate a fast and large postsynaptic response. Because the individual neuron’s response is close to all-or-none — a binary switch from quiescent to depolarized and spiking — these two mechanisms do not trade off against each other: the synaptic filter can be as stringent as needed for selectivity without compromising the speed or reliability of neurons that are successfully recruited. The decision to initiate escape then emerges at the dPAG population level: threat input stochastically recruits neurons across the dPAG network, with stronger threats switching on a larger fraction, and escape is initiated when this fraction is sufficiently large. This architecture also means that the system can be readily modulated by shifting the transition point of individual neurons — for example through changes in tonic inhibition ^28,29^ — which will change the fraction recruited for a given level of threat input, and therefore the escape threshold, without affecting the speed or fidelity of neurons that do respond.

A striking feature of dPAG neurons is that despite very high intrinsic excitability they do not fire during exploration, because the synaptic input rate is not high enough to reach the action potential threshold. This combination produces a system that is simultaneously quiet and primed to respond to even a modest increase in synaptic drive — here, approximately 25 Hz above baseline. This finding also implies that release probability at synapses onto dPAG neurons, or the activity of their presynaptic partners, must be tightly regulated during non-threatening conditions, since even small increases in synaptic drive would produce spontaneous escape-like responses. High excitability also serves a second function: it converts temporally irregular synaptic input into a sustained depolarization because the high input resistance prolongs the membrane time constant, which favours summation of discrete synaptic events. The synaptic drive during threat is not a smooth current but a stochastic stream of discrete events at modest rates, which in a less excitable neuron would produce noisy, intermittent depolarization and discontinuous escapes. The stable voltage step we observe provides the continuous drive required for sustained escape.

The mechanism we describe — threat detectors connected to highly excitable neurons that generate fast, reliable escape responses — is strikingly reminiscent of the giant fibre systems that mediate escape in invertebrates and fish ^30^. In crayfish, drosophila, and teleosts, large-calibre neurons with low spike thresholds receive direct input from threat-sensing circuits and trigger stereotyped escape actions with minimal delay ^31,32^. The functional logic is the same: a specialised neuron that is easy to activate and responds in an all-or-none manner, ensuring that once a threat is deemed genuine, the escape command is executed quickly and without question. A key difference, however, is that invertebrate systems typically rely on one or a small number of identifiable giant neurons, whereas the mammalian dPAG implements the same principle across a population of smaller neurons. This population architecture provides additional capabilities: the escape threshold can be modulated by controlling the fraction of recruited neurons rather than the properties of a single cell, and escape vigour can be graded at the population level while preserving quasi-binary responses in each individual neuron. The finding that the same core motif of high excitability appears across such distant lineages, from invertebrate giant fibres to mammalian midbrain populations, suggests that it may reflect convergent evolution of a common computational solution to the fundamental problem of escaping fast enough to survive.

## Methods

### Animals

All experiments were performed under the UK Animals (Scientific Procedures) Act of 1986 (PPLs 70/7652, PFE9BCE39 and PP2131611), following local ethical approval (Sainsbury Wellcome Centre, UCL).

Adult (6–12 weeks old), male and female C57BL/6J wild-type (Charles River), vGluT2-ires-Cre driver line (Jackson Laboratory, Stock No. 016963) and vGAT-ires-Cre driver line (Jackson Laboratory, Stock No. 028862) mice were singly housed with ad libitum access to water and chow on a 12 h reversed light cycle and tested during the dark phase. For freely moving single-unit recordings, mice were put under food restriction prior to behavioural recordings. Weight was checked daily and maintained above 85%.

### *In vivo* whole-cell recordings in head-fixed animals

#### Surgical procedures

##### Head-Plate Attachment

Animals were anaesthetised with isoflurane (3% for induction; 0.5–2% for maintenance, in oxygen at 1 L/min), and carprofen (5 mg/kg) was administered subcutaneously for perioperative analgesia. The fur on the forehead was shaved, the skin was disinfected, and eye gel was applied (Lubrithal). The animal was placed on a heating pad to maintain body temperature at 37°C and positioned in a stereotaxic frame (Kopf Instruments). An incision was made to expose the skull, and the edges were secured using tissue adhesive (Vetbond, 3M). The skull was cleaned of connective tissue, and a stainless-steel headplate (Neurotar) or an in-house custom-made headplate was positioned over the target area and affixed using light-cured dental cement (RelyX Unicem 2, 3M). Following surgery, animals were returned to their home cages and allowed to recover for a minimum of two days before further procedures.

##### Viral injection and fibre optic implantation for opto-tagging experiments

VGluT2-ires-Cre mice were anaesthetised and prepared for surgery as described above. A small craniotomy was made on the left hemisphere (AP: −0.2 mm, ML: 1.0 mm from Lambda), using 0.5 mm and 0.3 mm tungsten carbide burrs (Busch). Viral constructs (AAV2/2 EF1a-DIO-hChR2(H134R)-EYFP-WPRE) were injected using a pulled cut glass pipette (10 μL Wiretrol II, prepared with a Sutter P-1000) mounted on a stereotaxic injector (Janelia Research Campus, Ronal Tool Company), coupled to an oil hydraulic micromanipulator (MO-10, Narishige). The injection pipette was angled 30° from vertical to the left surface, and two injections were made at different depths (AP: −0.2 mm from Lambda, ML(30°): −0.63mm (left), DV(30°): −2.3 and −2.1 mm). A volume of 125 nL was delivered at a rate of ∼25 nL/min at each injection site. Following injection, the pipette was left in place for 10 minutes before slow retraction. A fibre optic cannula (200µm diameter fibre, numerical aperture 0.5, HealthiGlobal) was then inserted into the same craniotomy at a similar angle, with a slight offset (AP: −0.2 mm, ML: −1.07, DV: −1.45 mm), and affixed to the skull using light-cured dental cement. The head plate was then attached as described above.

##### Craniotomy for patch clamp recordings

Achieving sufficient stability for whole-cell recordings during exploration and escape episodes required extensive optimisation of the surgical procedure. On the day of recording, animals were anaesthetised with isoflurane (0.5–2% in oxygen at 1 L/min) and positioned in a stereotaxic frame (Kopf Instruments). A large circular craniotomy (∼2.5 mm diameter) was made over the right hemisphere using a handheld dental drill with 0.5 and 0.3 mm tungsten carbide burrs (Busch), targeting the area above the transverse and sagittal sinuses, centred at approximately −4.8 mm AP and 1.0 mm ML from Bregma to expose the superior colliculi. A ∼2 mm durotomy was performed using a 27G needle in the mediolateral direction, positioned as close as possible to the posterior edge of the transverse sinus. To maximise recording stability and access to the PAG, a custom-made implant was used (adapted from ^33^; Extended Data Figure 1). The implant consisted of a short stainless-steel tube (2.41 mm inner diameter, 1.78 mm inner diameter, 1 mm height; MicroGroup, Medway, MA) attached with superglue (Loctite) to a 2.6 mm diameter Kapton disc with seven 500 μm-diameter laser-cut holes (125 μm thick; Laser Micromachining Limited). The disc extended beyond the outer diameter of the tube, forming a lip that could be slid beneath the dura and sinus, stabilising against the skull edges of the craniotomy. The implant was positioned such that the rim of the tube could gently push the sinus anteriorly, while the implant applies mild downward pressure over the brain surface, with the holes providing access for electrode penetration to the PAG and SC. The implant was affixed to the skull with light-cured dental cement, and the exposed brain was covered with warmed 3% agarose in saline and sealed with silicone elastomer (Kwik-Cast, WPI). A pin to hold the ground was secured to the contralateral skull with dental cement. Following surgery, animals were returned to their home cage to recover for at least two hours before recordings began.

#### Head-fixation setup and data acquisition

##### Behavioural recordings

We used the Mobile HomeCage large system (Neurotar), which includes a carbon-fibre air-lifted arena (290 mm diameter) with a 23 mm transparent wall. Compressed air was delivered via a wall-mounted source at a pressure of 30 psi. To clamp mice to the head-fixation system, mice were briefly anaesthetised (<2 min) using isoflurane (2-5% in oxygen at 1 L/min) and rapidly connected to the head clamp. Mouse position was continuously tracked at 100 samples/s using the built-in magnetic locomotion tracking system, controlled by the Mobile HomeCage software (Neurotar). Mice were video-recorded from the side using a near-infrared GigE camera (Basler ace) at 50 frames/s. The setup was enclosed within a blackout frame lined with nylon light-blocking fabric (Thorlabs) and illuminated using infrared LEDs. Additional ambient illumination was provided by a monitor (Dell, 37 × 30 cm) positioned above the mouse at a 45° angle and 15° left offset, at a distance of 30 cm, producing ∼75 lux at the level of the mouse’s eyes. At the start of each recording session, a shelter made of bedding from the mouse’s home cage was placed along a 10 cm red acetate-shaded wall inside the arena.

##### Threat stimuli

The visual stimulus consisted of a sequence of three 1 s presentations of a 95% contrast, linearly expanding black circle (visual angle: 2.2°-168.2°, expansion rate: 166°/s), interleaved with 500 ms intervals. The auditory stimulus was a sequence of three 1 s 10 Hz waves, increasing in volume from 0 to 75 dB, also separated by 500 ms. In a subset of experiments, the sweeping auditory stimulus was used, consisting of a 3-s frequency-modulated upsweep from 17 to 20 kHz. Sound pressure level near the mouse’s head was ∼75 dB.

##### Synchronization

To ensure precise synchronization across all acquisition and stimulation channels, including video recording, locomotion, visual and auditory stimuli, and electrophysiological recordings, a custom LabVIEW program (Mantis, LabVIEW 2015, National Instruments (NI)) controlled four BNC-2090A NI boards, enabling all inputs and outputs to be aligned to a shared hardware clock. The software ran continuously during the whole session. Visual stimuli were generated using a custom MATLAB script using Psychtoolbox-3 ^34,35^. To account for monitor response delays, a photodiode (Thorlabs) was placed at the top-left corner of the screen. It was illuminated by a 1 × 1 cm black rectangle presented concurrently with each visual stimulus, and the photodiode signal was directly recorded by the NI board for precise timing alignment. Auditory stimuli were generated in Mantis and sent as analogue signals (300 kHz sampling rate) to the NI board, then routed through a Tucker-Davis Technologies amplifier to an L60 ultrasound speaker (Pettersson), positioned 50 cm above and in front of the mouse. Electrophysiological signals were acquired via Mantis by direct control of MultiClamp Commander software (Molecular Devices), with command signals delivered either directly from MultiClamp Commander or routed via Mantis and the NI system to the MultiClamp amplifier. Side-camera frames were triggered via a digital output from Mantis, and the camera trigger signal was simultaneously recorded as input. To validate synchronization between video and stimulus onset and to detect potential dropped frames, an infrared LED (LED405E, Thorlabs) within the camera field of view was pulsed concurrently with each visual or auditory stimulus. Signals from the locomotion tracking system were synchronised using a shared trigger between the Neurotar software and Mantis. Speed and XY position values were extracted from the Neurotar tracking data (100 Hz sampling rate) and aligned to the Mantis acquisition clock using this trigger signal. Values were then interpolated and smoothed to produce a continuous trace at the Mantis sampling rate (30 kHz).

#### Habituation

To habituate mice to head-fixation, training sessions lasting 1-1.5 h were conducted once daily for 3-5 days, depending on each mouse’s individual performance. Performance was evaluated based on the percentage of time spent moving within the arena and the total running distance. Mice were considered ready for recordings once their movement time exceeded 18%. In some cases, the first habituation session was performed in the Mobile HomeCage small system (18 cm diameter, 7 cm dark walls), placed within a triple-clamp setup that allowed simultaneous training of 2-3 mice.

#### *In vivo* recordings and opto-tagging

Whole-cell patch-clamp recordings followed the procedure described in ^36^, using a Multiclamp 700B amplifier (Molecular Device) and monitored using an HMO1002 oscilloscope (Rohde & Schwarz). Signals were sampled at 30 kHz and filtered at 10 kHz. Pipettes were pulled from borosilicate filamented glass capillaries (Harvard Apparatus, 1.5 mm OD, 0.85 mm ID, 75 mm L) with a PC-10 Narishige puller to a final resistance of ∼4–7 MΩ. Pipettes were backfilled with filtered internal solution containing (in mM): 130 KMeSO_3_, 10 KCl, 10 HEPES, 4 NaCl, 4 Na-phosphocreatine, 4 Mg-ATP, 1 EGTA, 0.5 Na_2_GTP, 285–290mOsm, pH was adjusted to 7.3 with KOH.

At the start of the session, mice were head-fixed to the head-clamp and the silicone elastomer and agarose covering the craniotomy were removed. The ground wire was connected to the pin and placed on the skull, in the vicinity of the craniotomy. In opto-tagging experiments, the fibre-optic patch cord was attached to the fibre implant tip. Mice were given 1-2 min to recover from the brief anaesthesia before recordings commenced. Recording pipettes were pressurised to 500-600 mBar (monitored with a Keyence Pressure Sensor) and directed vertically to one of the implant holes using the Junior in-vivo 4-axes micromanipulator (Luigs & Neumann). The pipette position was zeroed in the z-axis upon contact with the brain surface. Warmed 3% agarose in saline was then applied over the craniotomy to stabilise the pipette during recordings. The pipette was lowered to approximately 200 μm above the target recording depth (typically 1.2 mm below the brain surface for PAG recordings), and the pressure was carefully reduced to 40-80 mBar, taking care not to fully release it. To search for a cell, the pipette was advanced in 2 μm steps until a change in resistance was detected, at which point the pressure was released to obtain a gigaohm seal.

All cells were initially recorded in current-clamp mode to assess membrane responses to current injections of increasing magnitude (−40 to +200 pA, 20 pA increments, 250 ms duration, with the upper limit adjusted based on cell excitability) and to determine resting membrane potential. Voltage-clamp recordings were performed at a holding potential of −60 mV, except for IPSC recordings, which were performed at −70 mV. Access resistance was continuously monitored and corrected offline and only cells with stable resistance were included in the analysis. In opto-tagging experiments, ChR2 stimulation was delivered using a 473 nm laser module (Stradus, Vortran) with direct analogue modulation. Light was applied as continuous pulses of 0.1-2 s duration, with laser power set to 15–20 mW measured at the tip of the optic patch cord before the experiment.

#### Histology

##### Localisation of recorded cells

To determine the precise anatomical location of recorded cells, the recording pipette was retracted at the end of most experiments and loaded with a fluorescent dye (DiI, DiD, or DiO; Vybrant, Invitrogen) using a fine brush, before being reinserted to the original recording depth for 5-15 min. Following the recording session, brains were extracted and fixed in 4% paraformaldehyde (PFA) for a minimum of 16 h at 4°C, before transferred to PBS, for up to two weeks prior to processing. To confirm the fluorescent dye track and expression of ChR2, either a 3D whole-brain dataset was obtained using serial micro-optical sectioning 2-photon tomography ^37^, or 40-100 μm thick frontal sections were cut in 0.1 M PBS using a HM650V vibratome (Microm International), mounted using SlowFade Gold Antifade Mountant with DAPI (ThermoFisher) and imaged on a wide-field fluorescence microscope (AxioImager 2, Zeiss). Brains which were 3D imaged were aligned to the Enhanced and Unified Mouse Brain Atlas ^38^ using the BrainGlobe tool brainreg ^39–41^. This atlas contains annotations of the individual PAG subregions taken from the Franklin-Paxinos Mouse Brain Atlas ^42^. Dye tracks of patch pipettes were segmented in atlas space using the brainGlobe-segmentation tool. In cases where tissue damage prevented reliable atlas registration, tracks were manually segmented with brainreg based on visual comparison to histological images. The end point of a segmented track was taken as the location of a recorded cell. For cells which were recorded in very close proximity but were not marked by a dye track, their position was estimated along the segmented track trajectory. These estimations were based on the relative depths of unlabelled and dye labelled cell recordings and accounted for the maximal radius a pipette could have been inserted within, given the dimensions of the cranial implant which pipettes were inserted through. Cell coordinates were then visualized using brainrender ^43^. In all other brains, the position of the pipette tip was manually annotated to the Paxinos and Franklin Mouse brain atlas. Cells located in the dmPAG, ldPAG and lPAG were included in the analysis.

##### Confirmation of fibre placement and injection sites

For histological confirmation of fibre placement and injection sites in opto-tagging experiments, a 3D whole-brain dataset was acquired using the same procedure as described above.

#### Data analysis

##### Behaviour

Exploration in the Mobile HomeCage arena was defined as periods during which the mouse’s speed exceeded 10 mm/s. Distance to shelter was computed at each frame as the Euclidean distance between the animal’s tracked position and the shelter centre coordinates. Spontaneous speed traces were randomly selected for each experiment from epochs that did not include any stimulus presentation in the 40 s preceding or during the epoch. The proportion of time spent in the shelter was calculated by identifying all time points at which the mouse was located within a virtual shelter boundary, defined as a 60° angular sector covering the outer half of the arena radius, positioned at the top of the arena. The proportions of time spent exploring and of time spent in the shelter were calculated from 61 recording sessions, using only the period preceding the first stimulus presentation, and restricted to sessions in which the stimulus was introduced at least 30 min after session onset. Escape probability and reaction time were calculated from 21 behavioural experiments. Trials were defined as escapes when the animal’s speed exceeded 10 mm/s for a cumulative duration of at least 0.8 s within 5 s of stimulus onset. The remaining trials were classified as failures, or as startles if speed briefly exceeded 10 mm/s but for less than 0.8 s cumulatively. For analysis of escape vigour, escape trials were further subdivided into low-vigour escapes, in which speed never exceeded 70 mm/s within 7 s of stimulus onset, and high-vigour escapes comprising all remaining escape trials. This was followed by manual curation of speed traces and video recordings to correct for misclassifications. Escape reaction times were calculated as the first sample following stimulus onset, in which speed exceeded 10 mm/s.

##### Physiology

The resting membrane potential was determined by averaging the membrane voltage (Vm) over the first second of the recording. Resting Vm traces were mean subtracted to produce the Vm distributions in Fig. 1D. The action potential threshold was defined as the membrane voltage at which the phase plot (dV/dt vs. V) showed a sharp upward deflection, identified manually from the first action potential evoked by the lowest current step. Input resistance (Rin) was calculated from the steady-state voltage deflection in response to a hyperpolarising current pulse (−40 pA, 250 ms duration, 4-8 repetitions). The membrane time constant (τ) was estimated by fitting a single exponential to the voltage decay following the offset of a 40 pA current pulse. The difference between the resting membrane potential and the action potential threshold was calculated by subtracting the threshold from the resting potential. Rheobase was defined as the lowest current step (20 pA increments, 4-8 repetitions per step) that evoked at least one action potential across repetitions.

EPSPs were detected from current-clamp recordings using a deconvolution-based method ^44^. First, an EPSP template was built from a representative recording with low series resistance by averaging manually selected unitary events with clear, isolated waveforms. Each experimental trace was then deconvolved with the template in the frequency domain. The resulting deconvolved signal was filtered and histogram-fitted to estimate baseline noise σ. EPSPs were detected when local maxima of the signal exceeded a threshold of 4×σ, and reviewed manually to keep only events with a clearly identifiable unitary peak. All EPSPs from the same neuron were averaged to measure peak EPSP amplitude. EPSCs were detected from voltage-clamp recordings using a manually set threshold crossing on the first derivative, and all EPSCs from the same neuron were averaged to measure peak EPSC amplitude. The frequency of spontaneous EPSCs was computed from inter-event intervals (IEIs), and a probability density was then computed.

To visualize the membrane potential dynamics during escape, voltage traces from all recorded cells were aligned to stimulus onset and zeroed by baseline mean subtraction. For trial classification only, traces were z-scored over a window spanning 1.67 s before to 3 s after stimulus onset, and the first two principal components were extracted using PCA. Trials were then clustered into two groups using k-means clustering on the first two principal components. The cluster exhibiting stimulus-evoked depolarization was identified visually and selected for further analysis. The peak amplitude of the averaged response was calculated for the top 5% of voltage samples, during 1 s following stimulus onset. For the histograms shown in Fig. 2C, Vm traces were mean subtracted and then the top 5% of voltage values for each cell were computed for 1s before stimulus onset and for the stimulus duration, to generate Vm histograms for rest and threat periods, respectively.

To warp the responses to escape, voltage and speed traces were aligned to escape onset, linearly interpolated from escape onset to escape offset to the median escape duration across trials, and averaged. For each trial, a significance threshold was defined as the mean plus two standard deviations of the baseline voltage computed over the 333 ms window preceding stimulus onset, and voltage traces were binned into 20 ms bins. A bin was classified as significant if its mean voltage exceeded this threshold. Response onset was defined as the start of the first significant bin following stimulus onset. Voltage step onset in relation to escape onset was defined as the median of all onsets.

To examine the relationship between voltage step duration and escape duration, the same significance criterion was applied to stimulus-aligned traces from escape onset to escape offset. Only trials in which the mean voltage in the 670 ms window following stimulus onset exceeded the threshold were included. The total duration of significant bins was correlated with escape duration across trials using Pearson correlation. The spontaneous EPSC rate over time was computed in 150 ms bins, for escape trials only. To compare the fraction of active trials in different behavioural outcomes, voltage and speed traces were averaged separately for each outcome (failure, startle, low-vigour, high-vigour escape, and all escape). A trial was classified as significant if the mean voltage in the 667 ms window following stimulus onset exceeded the mean plus two standard deviations of the baseline voltage computed over the 333 ms window preceding stimulus onset. The fraction of significant trials was computed for each behavioural outcome as the number of significant trials divided by the total number of trials of that type.

To compare responses to visual and auditory threat stimuli, voltage traces from trials with significant depolarizing responses were used, and spikes were removed as described above. The mean squared difference between each voltage sample was computed for all pairwise trials, within threat type, and across threat types, for PAG and SC cells separately.

In optotagging experiments, a neuron was classified as directly activated by ChR2 if a brief light pulse evoked a robust step depolarisation of at least 10 mV within 5 ms of light onset. A depolarization response to threat was defined manually as an increase in membrane potential following stimulus onset.

### Single-unit recordings in freely moving animals

#### Surgical procedures

##### Neuropixels probes implantation

Five VGluT2-ires-Cre mice were used in this part of the study. Surgical procedures were performed as described above and previously ^23^. In a first surgery, animals received a viral injection of AAV2/2-EF1a-DIO-hChR2(H134R)-EYFP into the dorsal PAG. At least 8 weeks later, a second surgery was performed in which an optic fibre was implanted unilaterally above the dPAG at a 35° mediolateral angle. A 4-shank Neuropixels 2.0 probe was chronically implanted targeting PAG, SC, IC and PMd at a 40° anterior-posterior angle, and a second probe (single- or 4-shank Neuropixels 2.0) was implanted targeting ACC and VMH at a −35° anterior-posterior angle (data from the second probe were not included in this study). Probe shanks were coated with DiD (1 mM in ethanol, Invitrogen) prior to insertion for track identification.

#### Behavioural setup and data acquisition

##### Behavioural recordings

Behavioural experiments were performed in an elevated white, circular Perspex arena (92 cm diameter) with a red Perspex shelter (20 × 10 × 10 cm; 12 cm wide entrance) placed at the edge of the arena. The arena was housed within a dark, sound-deadening enclosure (140 x 140 x 160 cm) illuminated by 6 infrared LED lights (Abus). Experiments were recorded at 40 frames/s with an overhead near-IR GigE camera (acA1300-60gmNIR, Basler). Video recording and sensory stimulation acquired and synchronized using a PCIe-6351 and USB-6343 board (National Instruments) controlled with Bonsai software ^45^. A constant 1 Hz train generated by one of the boards was fed back to both behavioural boards and the Neuropixels board to synchronize the data streams.

##### Threat stimuli

A projector (BenQ) pointed at an overhead screen (Xerox) was used for visual stimuli and additionally provided a background luminance of 3 lux. The visual threat stimulus was a sequence of 5 expanding dark circles that expanded linearly over 400 ms, then maintained the same size for 250 ms and repeated with a 500 ms interstimulus interval. An overhead ultrasound speaker (L60, Pettersson) was used to deliver auditory threat at ∼80dB at the arena floor. The auditory threat stimulus was either a 9-s 10 kHz tone or a sequence of three 10 kHz tones whose amplitude increased linearly over 1 s and repeated with a 500 ms interstimulus interval.

#### Single unit recordings

A rotary joint (either AHRJ-OE_PT_AH_12_HDMI or AHRJ-OE_1×1_PT_FC_24_HDMI+4, Doric) was used to prevent twisting and torsion cables. Signals from the Neuropixels probes were recorded using SpikeGLX at a sampling rate of 30 kHz and high-pass filtering at 300 Hz. Active recording sites were chosen based on expected probe depths and a whole probe activity survey done prior to recording sessions.

At the end of the experiment, to confirm the location of probe implantation, the mouse was perfused and a 3D whole-brain dataset was acquired using the same procedure described above. The brain was registered to the Allen Brain Atlas using brainreg and tracks were manually reconstructed in brainreg-segment. Active sites were assigned to brain regions based on the registration and additionally adjusted based on electrophysiological signatures.

Neuropixels data were pre-processed using CatGT (https://github.com/billkarsh/CatGT) as previously described ^23^. Spike sorting was done using Kilosort 2.5 ^46^ and manually curated using Janelia Rocket Cluster 3.0 ^47^.

#### Data analysis

##### Behaviour

Behavioural videos and tracking data were sorted into peri-threat stimulus trials and manually annotated for escape occurrence, defined as a continuous movement consisting of a head orientation followed by full-body turn and run towards the shelter ^3^. Escape onsets were defined as the first video frame in which the mouse made the initial head orientation. Escape probability was calculated as the proportion of trials in a session where the stimulus triggered escape to the shelter. For each escape, reaction time was defined as the latency between stimulus onset and escape onset and maximum escape speed was defined as the peak speed reached between escape onset and shelter entry.

##### Neuronal responses

Firing rates of single units were aligned to stimulus onset and displayed as raster plots with 10 ms bins. To identify active units, spike rates were computed for each trial in 50 ms bins. A per-trial baseline firing rate was estimated from the 330 ms window preceding stimulus onset; if no spikes were present in this window, a 1-s pre-stimulus baseline was used instead, with a minimum threshold of 0.01 Hz. The significance threshold was defined as the mean plus two standard deviations of the baseline firing rate. A trial was classified as significant if the firing rate exceeded this threshold for at least four consecutive 50 ms bins (with a maximum gap of one bin allowed) within the test window. For escape trials, the test window spanned from stimulus onset to escape offset. For failure trials, the test window duration was matched to a fake offset drawn randomly from the escape offset distribution of the same recording session. Response onset was defined as the start of the first significant bin. A unit was classified as active, if it had at least one significant trial in either escape or failure. The response firing rate was computed in a 1 s window starting from the response onset of each trial, and the firing rate increase was defined as the difference between the response and baseline (1 s window before threat onset) rates. The fraction of significantly active units per trial was computed for each recording session across all active units, separately for escape and failure trials.

Spike times from significantly responding trials of active units were aligned to escape onset and warped for the escape period (escape onset to escape offset) by linearly interpolating to the median escape duration across trials. Warped PSTHs were computed in 10 ms bins and averaged across trials. Speed traces were similarly warped and averaged. The increase in firing rate per trial was computed as the difference between the mean firing rate during the escape period and the mean firing rate during a 1 s window before threat onset. For comparing escape and failure conditions, stimulus-aligned mean PSTHs were computed separately for significant and non-significant trials, for both escape and failure conditions. The firing rate change per trial was computed as the difference between the mean firing rate in the 1 s following stimulus onset and the mean firing rate at baseline.

#### General statistical analysis

Unless otherwise specified, all values are reported as mean ± s.d. Statistical significance was considered when P < 0.05. Data were tested for normality using the Shapiro–Wilk test, and a parametric test was used if the data were normally distributed, and a non-parametric test otherwise, as detailed in the text next to each comparison.

## Data availability

Data and code will be shared at the time of publication and are available before that upon request.

## Acknowledgements

This work was funded by a Wellcome Senior Research Fellowship (214352/Z/18/Z), by the Sainsbury Wellcome Centre Core Grant from the Gatsby Charitable Foundation and Wellcome (GAT3755 and 219627/Z/19/Z) and by a European Research Council grant (Consolidator #864912) (T.B.), by a Marie Skłodowska-Curie Individual Fellowship (706136), by an EMBO Long Term Fellowship (ALTF 879-2015), and by a Royal Society Dorothy Hodgkin Fellowship (DHF\R1\191089) (Y.L.), by a A*STAR National Science Scholarship (PhD; Y.L.T) and the SWC PhD Programme (Y.L.T.). We thank members of the Branco lab and T. Margrie for discussions and comments on the manuscript; We thank M. Velez-Fort and T. Margrie for initial guidance on *in vivo* whole-cell recordings; the SWC Neurobiological Research Facility and FabLabs for technical support; K. Betsios for programming the data acquisition software.

## Author contributions

T.B. and Y.L. conceived the project, designed the experiments, analysed the data and wrote the manuscript. Y.L. performed whole-cell recordings. Y.L.T. performed single-unit experiments; Y.L., G.F., Y.W. and A.F prepared and trained mice for head fixation, performed and tagged behavioural experiments and histology. A.F. Injected mice for optotagging experiments. M.H registered brains and located cells.

---

**Extended Data Figure 1:**
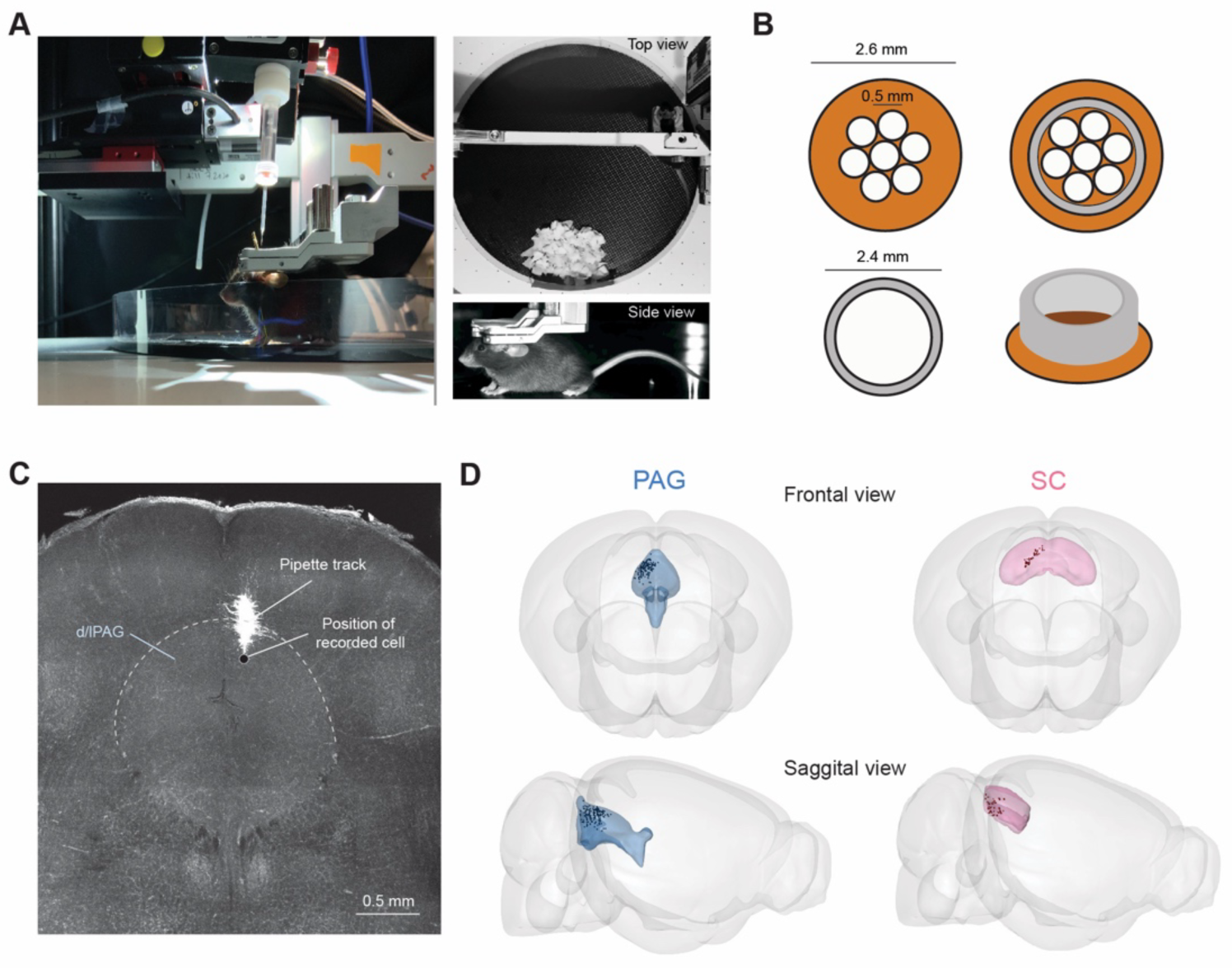
Head-fixed setup for whole-cell patch-clamp recordings during escape. **A,** Left, side view of the head-fixed setup showing the floating arena, head-fixation apparatus and intrumentation for whole-cell recordings. Right, top and side views of a head-fixed mouse. **B,** lllustation of the implant used for stabilising the brain during recording, consisting of a Kapton disc with seven holes (top left) and a short stainless-steel tube (bottom left) that were attached together (right, top and side views). The disc extended beyond the outer diameter of the tube, forming a lip that could be slid beneath the dura, to gently push the venus sinus anteriorly and apply a mild downward pressure over the brain, C, Example image of a Oil fluorescent track from a patch-clamp pipette, recovered after recording from a dPAG cell. **D,** frontal and sagittal views showing the positions of all recorded cells (black dots) in PAG (blue, n=163) and SC (pink, n=56)

**Extended Data Figure 2:**
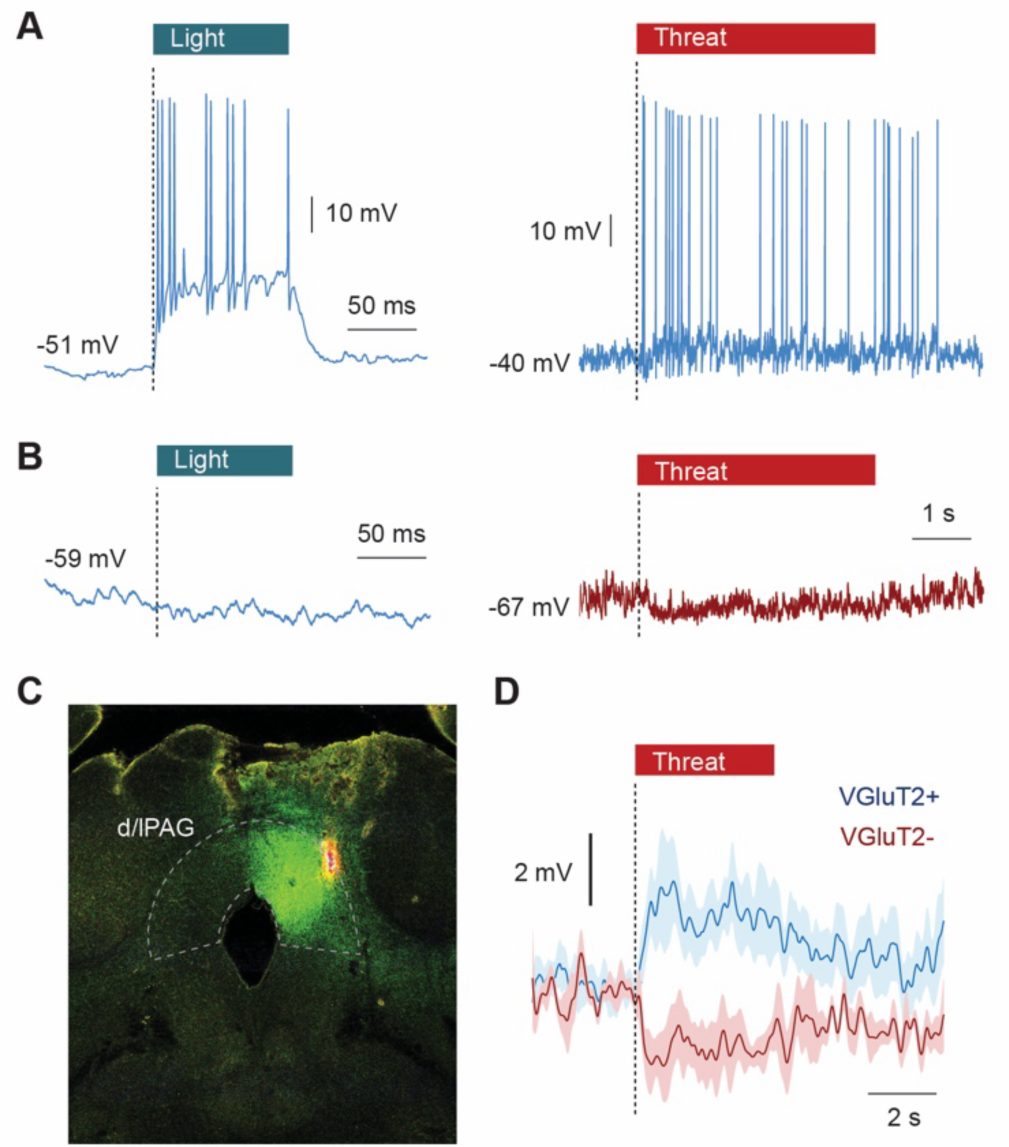
Optotagging VGluT2+ neurons in dPAG. **A,** Left, example voltage response to a 100 ms light pulse, indicating that the recorded dPAG cell was expressing ChR2 (whole-cell recordings were established blindly in VGluT2-Cre mice injected with DIO-hChR2-EYFP). Right, voltage response of the same cell to a sound threat, showing a depolarizing step response with action potentials. **B,** Left, example of voltage response to 100 ms light pulse in a PAG cell that did not express ChR2. Right, voltage response of the same cell to a visual threat, showing a hyperpolarizing response. C, Example frontal brain slice image of a Vglut2-Cre mouse expressing ChR2-EYFP in the PAG, and the lip of the fluorescence pipette track after a recording inm the dlPAG (Oil dye). **D,** Average voltage responses to threat from cells expressing ChR2 in VGluT2 cells (blue), and of cells that did not express ChR2, but were recorded in an area with high ChR2 expression (red).

**Extended Data Figure 3:**
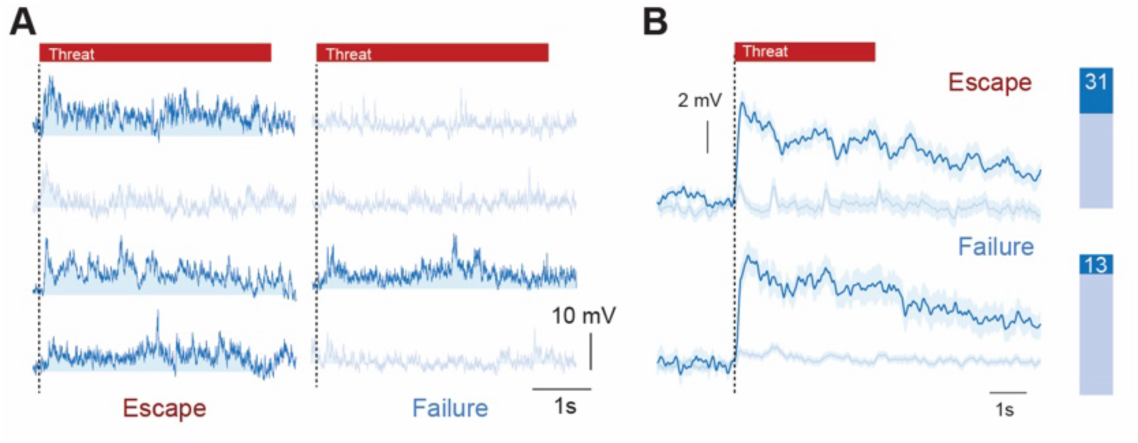
Escape and failure trials recorded in the same cell. **A,** Examples of voltage traces during escape (left) and failure (right) trials, from cells that had both trials recorded. Trials with a sigificant depolarizing response are coloured in darker blue. Each row shows a different cell. **B,** Average voltage traces for escape (upper) and failure (bottom) trials from all depolarizing (dark blue) and non-depolarizing (light blue) trials. Bars show the proprotion of each trial type for escape (top) and failure (bottom)

