## Extended data figures 1-3 for "A cellular midbrain mechanism for executing fast and reliable escape"

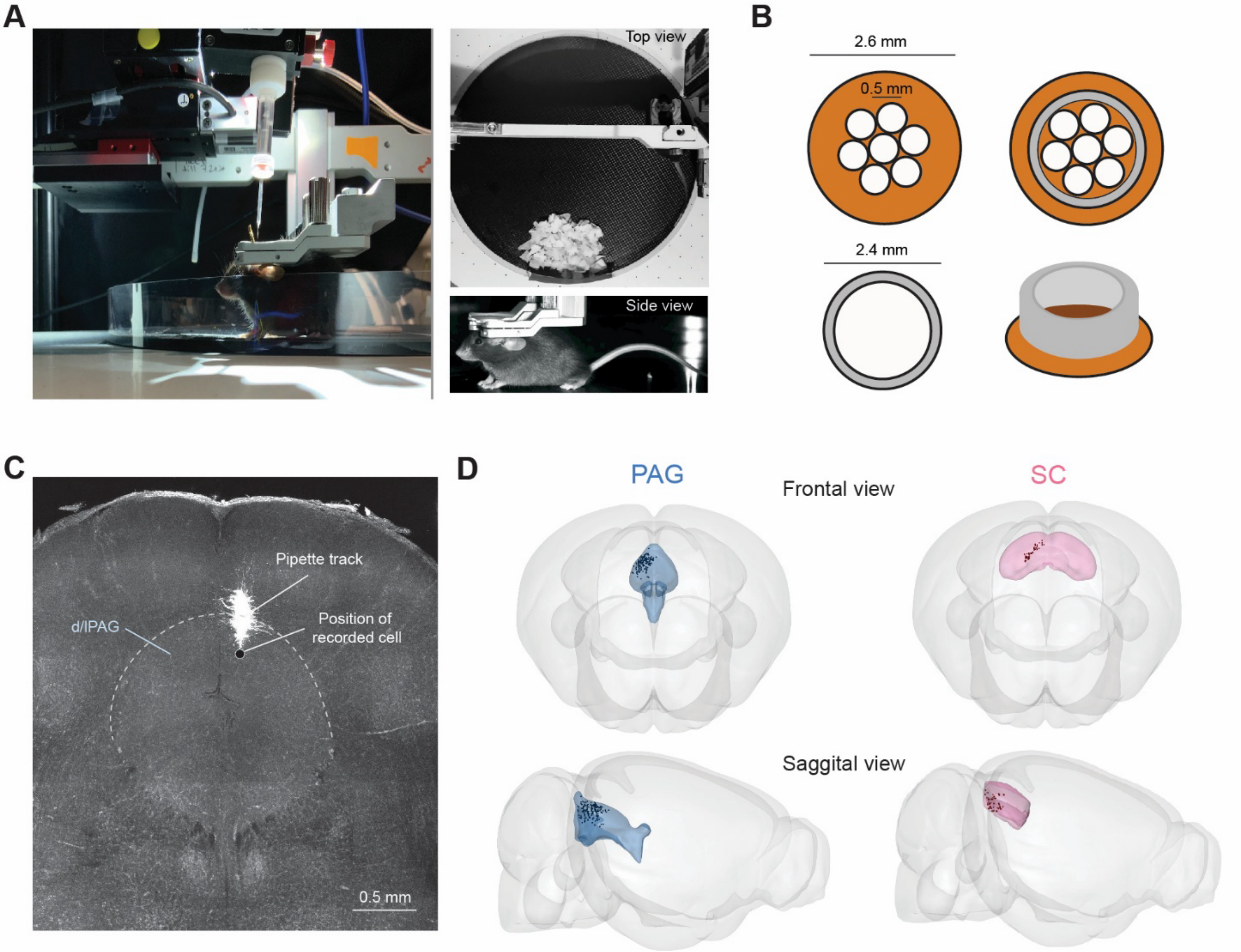


**Extended Data Figure 1: Head-fixed setup for whole-cell patch-clamp recordings during escape**

**A**, Left, side view of the head-fixed setup showing the floating arena, head-fixation apparatus and intrumentation for whole-cell recordings. Right, top and side views of a head-fixed mouse. **B**, Illustation of the implant used for stabilising the brain during recording, consisting of a Kapton disc with seven holes (top left) and a short stainless-steel tube (bottom left) that were attached together (right, top and side views). The disc extended beyond the outer diameter of the tube, forming a lip that could be slid beneath the dura, to gently push the venus sinus anteriorly and apply a mild downward pressure over the brain. **C**, Example image of a DiI fluorescent track from a patch-clamp pipette, recovered after recording from a dPAG cell. **D**, frontal and sagittal views showing the positions of all recorded cells (black dots) in PAG (blue, n=163) and SC (pink, n=56).

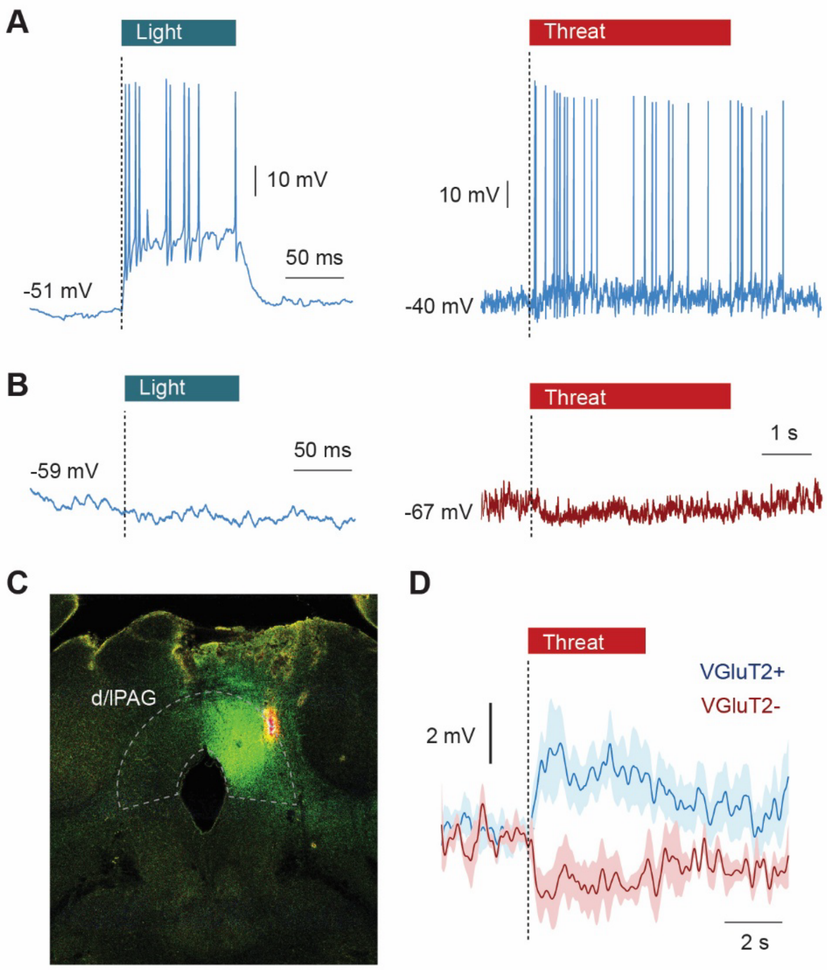


**Extended Data Figure 2: Optotagging VGluT2+ neurons in dPAG**

**A**, Left, example voltage response to a 100 ms light pulse, indicating that the recorded dPAG cell was expressing ChR2 (whole-cell recordings were established blindly in VGluT2-Cre mice injected with DIO-hChR2-EYFP). Right, voltage response of the same cell to a sound threat, showing a depolarizing step response with action potentials. **B**, Left, example of voltage response to 100 ms light pulse in a PAG cell that did not express ChR2. Right, voltage response of the same cell to a visual threat, showing a hyperpolarizing response. **C**, Example frontal brain slice image of a Vglut2-Cre mouse expressing ChR2-EYFP in the PAG, and the tip of the fluorescence pipette track after a recording inm the dlPAG (DiI dye). **D**, Average voltage responses to threat from cells expressing ChR2 in VGluT2 cells (blue), and of cells that did not express ChR2, but were recorded in an area with high ChR2 expression (red).

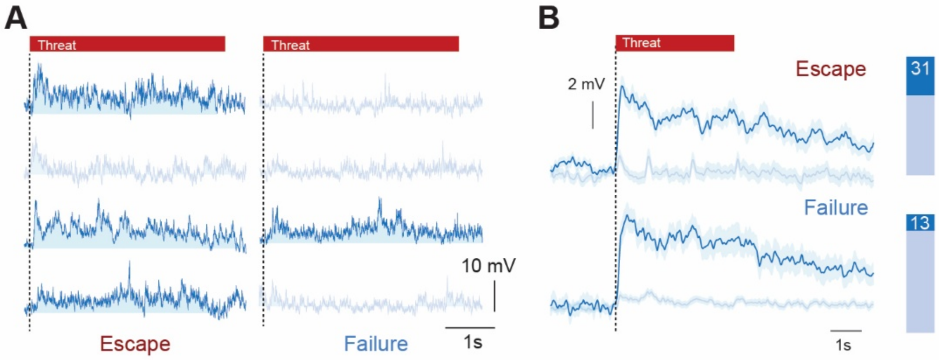


**Extended Data Figure 3: Escape and failure trials recorded in the same cell**

**A**, Examples of voltage traces during escape (left) and failure (right) trials, from cells that had both trials recorded. Trials with a sigificant depolarizing response are coloured in darker blue. Each row shows a different cell. **B**, Average voltage traces for escape (upper) and failure (bottom) trials from all depolarizing (dark blue) and non-depolarizing (light blue) trials. Bars show the proprotion of each trial type for escape (top) and failure (bottom)
